# Overcoming Observation Bias for Cancer Progression Modeling

**DOI:** 10.1101/2023.12.03.569824

**Authors:** Rudolf Schill, Maren Klever, Andreas Lösch, Y. Linda Hu, Stefan Vocht, Kevin Rupp, Lars Grasedyck, Rainer Spang, Niko Beerenwinkel

## Abstract

Cancers evolve by accumulating genetic alterations, such as mutations and copy number changes. The chronological order of these events is important for understanding the disease, but not directly observable from cross-sectional genomic data. Cancer progression models (CPMs), such as Mutual Hazard Networks (MHNs), reconstruct the progression dynamics of tumors by learning a network of causal interactions between genetic events from their co-occurrence patterns. However, current CPMs fail to include effects of genetic events on the observation of the tumor itself and assume that observation occurs independently of all genetic events. Since a dataset contains by definition only tumors at their moment of observation, neglecting any causal effects on this event leads to the “conditioning on a collider” bias: Events that make the tumor more likely to be observed appear anti-correlated, which results in spurious suppressive effects or masks promoting effects among genetic events. Here, we extend MHNs by modeling effects from genetic progression events on the observation event, thereby correcting for the collider bias. We derive an efficient tensor formula for the likelihood function and learn two models on somatic mutation datasets from the MSK-IMPACT study. In colon adenocarcinoma, we find a strong effect on observation by mutations in TP53, and in lung adenocarcinoma by mutations in EGFR. Compared to classical MHNs, this explains away many spurious suppressive interactions and uncovers several promoting effects.

The data, code, and results are available at https://github.com/cbg-ethz/ObservationMHN.

## 1 Introduction

Cancer progression models (CPMs) aim to describe and reproduce the evolutionary development of a healthy tissue into a malignant tumor, driven by a series of genetic (or epigenetic) events such as mutations or copy number alterations [4]. While these events occur randomly due to cellular replication errors, their establishment in the tumor cell population is not entirely random. It depends on the selective advantage they confer in the given environment and the genetic background [37]. Fixation of a genetic alteration in the tumor is often enabled or suppressed by previous events, making some chronological sequences and patterns of events more likely than others [43].

For instance, an initial mutation might promote tumor growth until it is starved for oxygen, whereupon subsequent mutations become beneficial that facilitate blood vessel formation. Access to blood vessels in turn sets the stage for further events culminating in metastasis. Conversely, some events can also suppress one another. This can result, e.g., from synthetic lethality, where some events aid the tumor cell individually but become fatal when they occur together. Or, events may target genes in the same regulatory pathway; whichever event occurs first disrupts the whole pathway, reducing selective pressure on the other event.

Such interactions between events are still poorly characterized, and learning them from data is the goal of CPMs. The challenge lies in the inherent limitations of available data. Datasets with many patients typically provide only bulk genotypes and do not resolve clonal structures. Most datasets are also cross-sectional: They provide a snapshot of many different tumors at a single time point each, but do not track tumors over multiple stages of their evolution.

While we do not know the time at which a tumor was observed relative to the start of its progression, it is also not entirely random: Tumors are usually detected rather late in this process, such that most data comes from later stages of the progression. Up to this point, the tumor has already grown, undergone changes and accumulated events. Some of these events have actually caused the tumor to grow. Thus, our ability to observe and study tumors depends on the events that occurred before we could detect the tumor. This dependence introduces a systematic bias to cancer progression models, which we resolve in this paper.

So far, CPMs have been developed that can be learned from bulk genotypes [16] but assume that tumors were observed at a random time independent of their progression events. They are trained on the co-occurrence patterns of events and model the probabilities of future events as functions of events already present. These functions define a causal network. CPMs build on the seminal work of Fearon and Vogelstein [18] who manually inferred from genetic and clinical data that colorectal cancer tends to progress along a chain of mutations in the genes APC→ KRAS→ TP53.

Oncogenetic Trees [15,3] extend such chains and allow each event to be a necessary precursor to more than one successor event. In Conjunctive Bayesian Networks (CBNs) [2,20,40] events may also require multiple precursors, thus extending trees to directed acyclic graphs (DAGs). CAPRESE [35] and CAPRI [46] are similar tree and DAG models where precursor events are not strictly necessary for successor events but raise their probabilities. Other DAG models with different functional forms are Disjunctive Bayesian Networks [42], Monotone Bayesian Networks [17] and Bayesian Mutation Landscapes [38]. Pathway Linear Progression Models [47] infer groups of mutually exclusive events and arrange them in a chain. PathTiMEx [14] generalizes this to CBNs of groups of mutually exclusive events. Network Aberration Models (NAMs) [28] are cyclic causal networks with promoting effects. HyperTraPS [31,24] and Mutual Hazard Networks (MHNs) [50] generalize this to cyclic networks with promoting and suppressive effects. Similar approaches [1,39] allow higher-order rather than pairwise interactions between events. TreeMHN [36] infers MHNs from intra-tumor phylogenetic trees derived from single-cell, multi-region or bulk sequencing data.

Here we address a fundamental oversight in all these CPMs: They do not include the observation of the tumor itself into their causal networks. Instead, we regard observation of the tumor as an event indicating that the tumor was biopsied, sequenced, and eventually included into the dataset. It implies that the tumor has become conspicuous due to its size, morphology, or symptoms such as weight loss, fatigue or pain. Since a dataset contains by definition only tumors at their moment of observation, neglecting any causal effects on this event makes CPMs prone to the notorious “conditioning on a collider” bias [27]. This bias is also known as Berkson’s paradox and refers to spurious associations between any variables that affect another conditioned variable. Joseph Berkson originally described it for a hospital in-patient population which showed a negative association between diabetes and cholecystitis [5], see Fig. 1. However, since diabetes is known to increase the risk for cholecystitis [12], one would naturally expect a positive association. The explanation for the spurious negative association is that diabetes and cholecystitis are both separate causes to be in the hospital and therefore to be observed in this study. Learning of one cause explains away the need for the other cause.

**Fig. 1.**
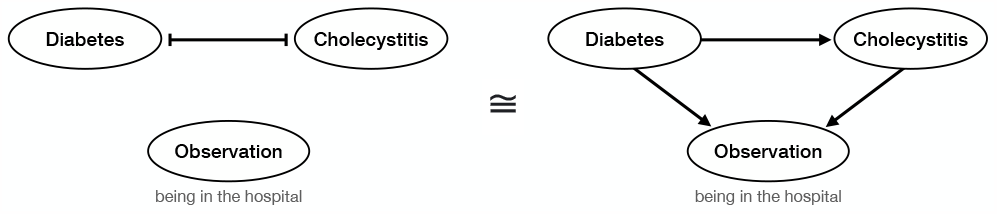
Berkson’s original example of a collider bias [5]: The negative association between diabetes and cholecystitis observed in hospital patients could be spuriously explained (left) by suppressive effects between the diseases, if the inclusion of a patient in the dataset were independent of both diseases. Alternatively, the same negative association can be correctly explained (right) by taking into account that both diseases have a promoting effect on being in the hospital, and thus observed in the dataset. The spurious association masks the actual promoting effect of diabetes on cholecystitis.

Similarly, the inference of CPMs from statistical associations can be grossly distorted when the observation should be a part of the causal network but is not properly accounted for. CBNs and MHNs are models in continuous time which do have a random observation event, but its rate is fixed at 1 and cannot be affected by other events. Timed Hazard Networks [11] extend MHNs by hidden variables for the observation times of all tumors, but these are also not affected by other events. NAMs [28] have an observation event whose rate depends on the total number of events that have occurred, but not on which particular events have occurred.

In this paper, we extend Mutual Hazard Networks by causal effects between its progression events and its observation event. Each event occurs at its base rate and has multiplicative effects on the rate of every other event. These effects can be greater than 1 (promoting), less than 1 (suppressive), or equal to 1 (neutral) and define a causal network with cycles. An MHN is a generative model of cancer progression in the form of a continuous-time Markov chain. We provide an analytical formula for its probability distribution over tumor states, explicitly conditioned on their times of observation. This formula uses tensor expressions which allows us to efficiently infer base rates and multiplicative effects between events via maximum likelihood estimation.

We demonstrate our approach on two datasets of colon adenocarcinoma (COAD) and lung adenocarcinoma (LUAD) from the MSK-IMPACT study [41,51]. Compared to classical MHNs, we find results that offer drastically different interpretations. In COAD, we find that TP53 strongly promotes observation, which explains away suppressive interactions and uncovers promoting effects between APC and TP53. For LUAD, the new model identifies EGFR mutations as principal observation drivers, which explains away its suppressive interactions with most other events but retains suppressive effects with KRAS.

## 2 Methods

We first summarize the definition of classical Mutual Hazard Networks from [50]. Then we extend MHNs by effects on the observation and derive a formula for their likelihood function. Finally, we show that such models are not uniquely identifiable from cross-sectional data and resolve this by a regularization that favors parsimony.

### 2.1 Classical MHNs with unaffected observation

Mutual Hazard Networks (MHN) [50] model cancer progression as a continuous-time Markov chain that describes how a tumor accumulates *n* possible progression events. Over the course of its progression, a tumor can be in any of 2^*n*^ states **x** ∈ {0, 1}^*n*^ where **x**_*i*_ = 0 encodes that event *i* ∈ {1,…, *n*} has not yet occurred and **x**_*i*_ = 1 that it has. We assume that every tumor starts at time *t* = 0 in the healthy state (0, …, 0)^⊤^ ∈ {0, 1} ^*n*^, accumulates events irreversibly one after another, and is finally observed at a random time *t* which is unknown.

Let **p**(*t*) be a vector of size 2^*n*^ that denotes the transient probability distribution over states at time *t* ≥ 0. Here we use a lexicographic order on {0, 1}^*n*^ with the leftmost bit cycling fastest, see Fig. 2 (bottom left). An entry **p**(*t*)_x_ denotes the probability that a tumor is in state **x** at time *t* ≥ 0. The initial distribution

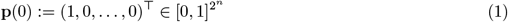

is concentrated on the healthy state. Its change over time is governed by the Kolmogorov forward equation:

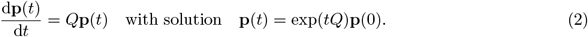

**Fig. 2.**
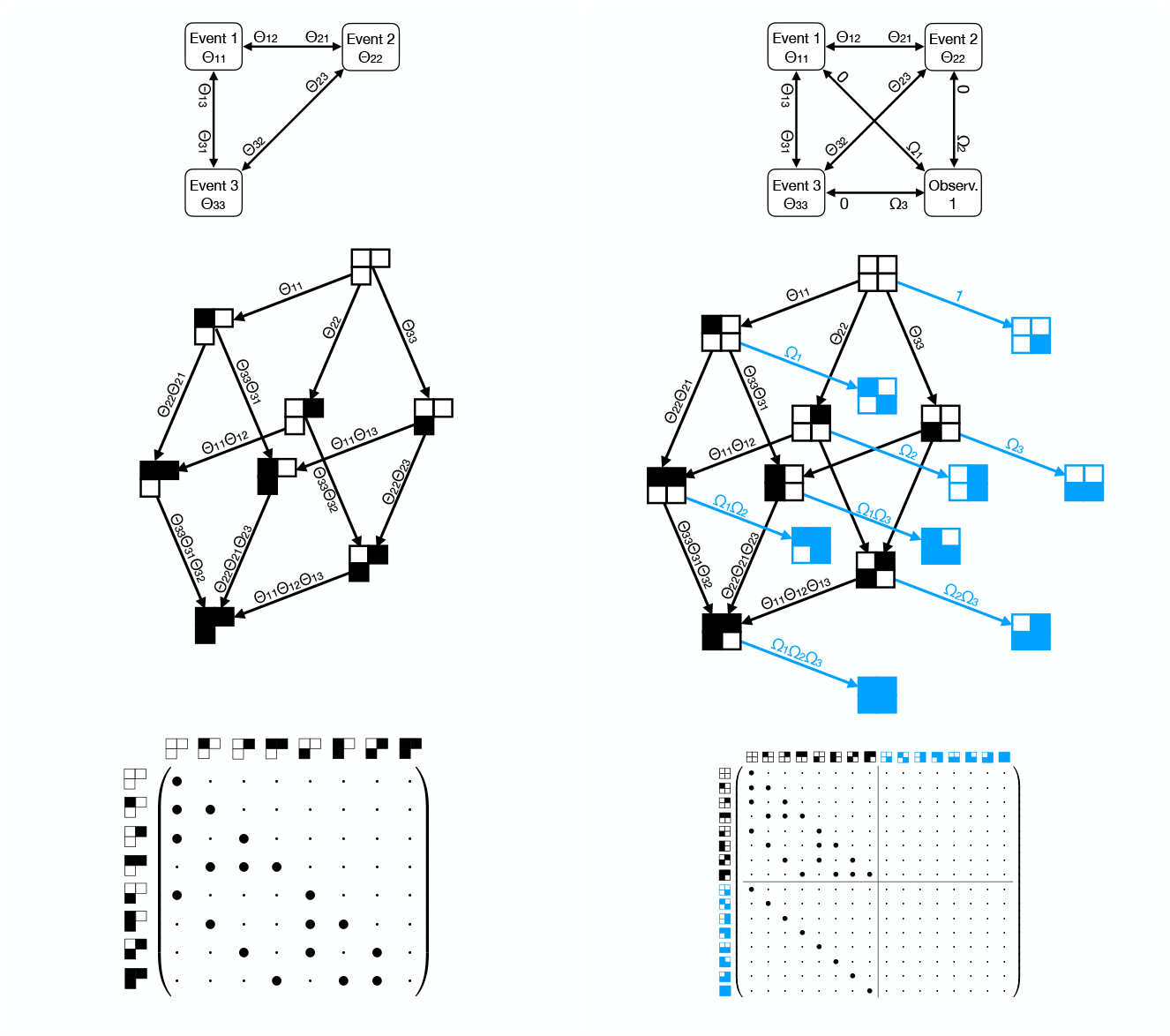
Comparative illustration for *n* = 3 progression events. Left: A cMHN with parameters *Θ* and implicit observation event whose rate is fixed at 1. Right: An oMHN with the observation as an explicit fourth event and parameters (*Θ*, Ω). For the cMHN and oMHN each: top: the corresponding causal interaction networks between events; middle: the transition rates of the corresponding Markov chains; bottom: the structure of the corresponding transition rate matrices for a lexicographic order of the state space.

Here, 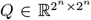 is the transition rate matrix, where an off-diagonal entry 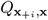 is the transition rate from a state **x** = (…, **x**_*i*−1_, 0, **x**_*i*+1_, …)^⊤^, which lacks event *i*, to the state **x**_+*i*_ := (…, **x**_*i*−1_, 1, **x**_*i*+1_, …)^⊤^, which differs from **x** only in the additional event *i*. By assumption, events accumulate irreversibly one at a time, and thus all other off-diagonal entries are 0 and *Q* is lower-triangular. Its diagonal entries are defined such that each column sums to 0.

Our aim is to learn for each event *i* how its rate depends on already present events in **x**. To this end, an MHN with parameters *Θ* ∈ ℝ^*n×n*^ defines the functional form

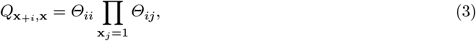

where *Θ*_*ii*_ > 0 is the base rate of event *i* and *Θ*_*ij*_ > 0 is the multiplicative effect of event *j* on the rate of event *i*.

In order to learn *Θ* from data via maximum likelihood estimation, we have to compute the probability distribution over all possible tumor states at the time of their observation. The observation in a classical MHN occurs randomly at a time which is exponentially distributed with a fixed rate of 1. Marginalizing over the unknown observation time *t* ∼ Exp(1) yields the time-marginal distribution

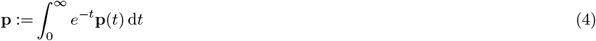

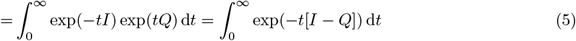

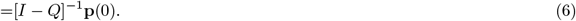

Note that eq. (5) is only valid for a fixed observation rate, since it relies on the fact that *Q* commutes with the identity matrix *I*. The log-likelihood of *Θ* for a dataset 𝒟 of observed tumor states is then

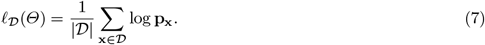

Maximizing the log-likelihood of *Θ*, e.g. via gradient ascent or quasi-Newton methods, requires operations that involve the huge matrix *Q*. To this end, we make use of the following representation of *Q* as a sum of tensor products:

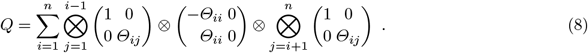

Using efficient tensor operations [9], *Θ* can be learned with a time and storage complexity only exponential in the number of events that have occurred for each tumor, rather than exponential in 2*n* [49].

### 2.2 MHNs with effects on observation

Here, we extend the classical MHN (cMHN) introduced in the previous section to an observation MHN (oMHN). We include the observation event into its causal network as an explicit (*n* + 1)^th^ event. Its base rate is defined as 1 in order to standardize the time scale. We introduce additional parameters Ω ∈ R^*n*^, where Ω_*j*_ > 0 is the multiplicative effect of the progression event *j* ∈{1, …, *n*} on the rate of observation.

Because now the observation rate depends on the state, we can no longer use eq. (6) to compute the probabilities of tumor states at their time of observation. Instead we use the following construction: We set the outgoing effects of the observation on all genetic progression events to 0. Once the observation occurs, it prevents all other events from ever occurring by multiplying their rates with 0, which freezes the data generating process at the time of observation^4^. The probability distribution at observation now equals the stationary distribution at infinity, which can be computed as follows.

Formally, we define the extended Markov chain on the state space

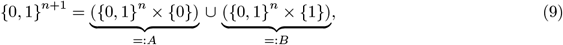

where *A* includes all states before observation and *B* all states after observation. The extended transition rate matrix 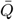 is of size 2^*n*+1^ *×* 2^*n*+1^ and has the following block structure:

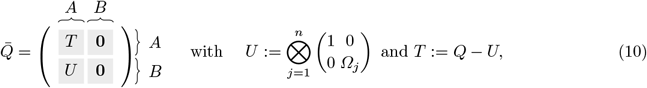

where each block is of size 2^*n*^ *×* 2^*n*^, see Fig. 2 (bottom right). The block *U* contains all transitions that introduce the observation event. It is diagonal with strictly positive eigenvalues and hence invertible. The block *T* contains all transitions that introduce a progression event, given by *Q* in eq. (8), and *U* is subtracted from its diagonal so that each column of 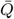 sums to 0. *T* is lower-triangular with strictly negative eigenvalues and hence also invertible.

The transient distribution 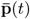 of the extended Markov chain can also be organized by blocks and is governed by the Kolmogorov forward equation:

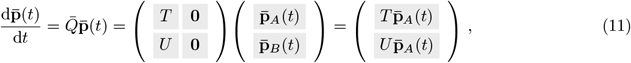

where 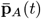 and 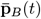 are each of size 2^*n*^ and denote the transient distribution restricted to *A* and *B* respectively. Given the initial distribution 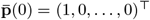, i.e.,

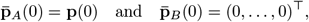

the solution to the Kolmogorov equation reads

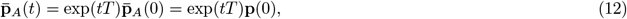

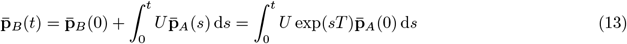

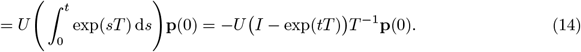

Because all eigenvalues of *T* are strictly negative, we can calculate the stationary distribution by

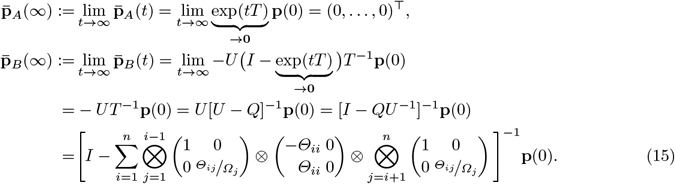

The log-likelihood of a dataset 𝒟 of observed tumor states is then

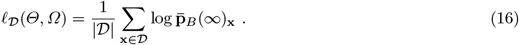

Computing and maximizing the log-likelihood to learn *Θ* and Ω has the same complexity as for a cMHN, i.e., it is exponential in the number of events that have occurred for each tumor.

### 2.3 Non-identifiability and regularization

Note that the formula for the stationary distribution of an oMHN (15) is the same as the formula for the time-marginal distribution of a cMHN (6) where the parameters *Θ*_*ij*_ are replaced by the fractions *Θ*_*ij*_*/*Ω_*j*_. It follows that an oMHN is not uniquely identifiable from cross-sectional data alone. For any oMHN with parameters *Θ* and Ω, we can construct a likelihood-equivalent cMHN with 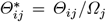 for *i* ≠ *j* and 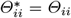, see Fig. 3. Although both models generate exactly the same observational data, they have very different causal interpretations. That is, if we intervened experimentally on the system, the two models would then differ in their future dynamics.

**Fig. 3.**
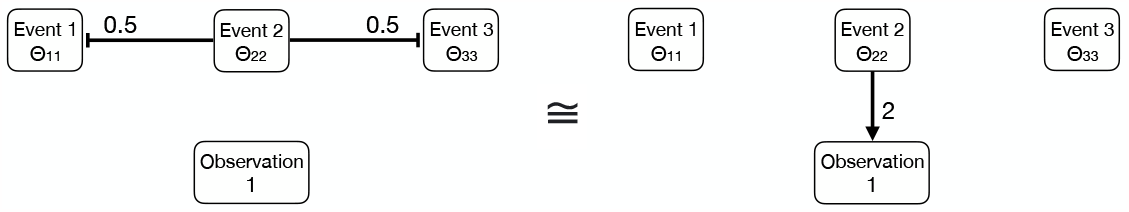
Example of a cMHN (left) and an oMHN (right) which generate the same observational data but differ in their causal interpretation. They imply different experimental predictions: A drug treatment which suppresses event 2 would increase the probabilities of events 1 and 3 according to the left model, but not according to the right model. (Both networks are fully connected, but neutral effects of multiplicative strength 1 are not drawn.)

In order to decide on a particular causal model, we cannot rely on data alone but have to incorporate background knowledge or preferences in the form of a Bayesian prior or a penalty on the likelihood. Following the principle of parsimony (Occam’s razor), we prefer simple models that postulate the least number of causal mechanisms for explaining the data. This means that MHNs should be sparse, in the sense that many effects *Θ*_*ij*_ = 1 and Ω_*j*_ = 1 for *i* ≠ *j*. In the example of Fig. 3, we would hence prefer the model on the right.

Moreover, we prefer symmetric models where many effects *Θ*_*ij*_ = *Θ*_*ji*_ since these are likely due to a single causal mechanism that is inherently symmetric, such as synthetic lethality or functional equivalence among mutations. While such effects presumably do not vary in strength whether event *i* or *j* occurs first, there may be important exceptions [45,29]. Hence, we do not want to impose strict symmetry on *Θ*.

To this end, we propose maximizing the log-likelihood regularized by the following penalty which induces sparsity and soft symmetry:

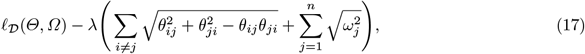

where *θ*_*ij*_ := log(*Θ*_*ij*_), *ω*_*j*_ := log(Ω_*j*_) and *λ >* 0 is a hyperparameter. Similar to the Group Lasso [56], this penalty promotes sparsity such that many logarithmic effects are 0 (hence multiplicative effects are 1) but pairs of effects *θ*_*ij*_ and *θ*_*ji*_ are selected together. The additional term −*θ*_*ij*_*θ*_*ji*_ ensures that symmetric effects of equal strength and sign are penalized only as strongly as a single effect *θ*_*ij*_ with *θ*_*ji*_ = 0.

In this paper, we choose *λ* in 5-fold cross-validation according to the One Standard Error Rule [26], which selects the largest value for *λ* such that its average log-likelihood is within one standard error of the optimum. We use this rule because the optimal *λ* tends to 0 for larger datasets as it becomes less necessary to prevent overfitting, but we still want to favor simple models to mitigate non-identifiability.

## 3 Results

We provide a new version of Mutual Hazard Networks with a corresponding efficient learning algorithm. These models shed new light on cancer progression by telling us which genetic events are most responsible for the clinical observation of a tumor. The models are thereby corrected for a collider bias and offer more realistic interpretations of cancer progression, showing fewer spurious interactions and more genuine interactions that had been previously overlooked.

Specifically, we applied our method and learned two models from somatic mutation data of colon adenocarcinoma (COAD) and lung adenocarcinoma (LUAD), which were originally collected by the Memorial Sloan Kettering Cancer Center [41] and retrieved through AACR GENIE [51]. We selected one primary tumor sample for each of the 2269 COAD patients and 3662 LUAD patients. As mutational events, we considered only likely pathogenic variants and selected the 12 most commonly affected genes in each of COAD and LUAD, as described in Supplementary S1.

Although oMHN models are more realistic than cMHNs, they are in principle equally powerful for explaining the data, so we did not necessarily expect results with higher likelihood. Nevertheless, we validated their model fit by splitting each dataset in half into a training and test set. We trained a cMHN and an oMHN on the training set and evaluated their log-likelihoods on the test set. For COAD, the cMHN achieved a log-likelihood of −5.14 while the oMHN achieved a slightly better −5.10. For reference, the independence model^5^ achieved −6.02 and the best possible performance was the entropy −4.48 of the test set. For LUAD, the cMHN achieved a log-likelihood of −3.96 and the oMHN achieved a slightly better −3.94. The independence model achieved −4.50 and the entropy of the test set was −3.74.

In the following, we report the models trained on the full datasets.

### 3.1 Colon Adenocarcinoma

Mutations in APC, KRAS and TP53 are long thought to be the cornerstones of conventional COAD progression [18]. The three events are abundant in the dataset (42%-72%) but enriched in samples with few events overall, see Supplementary S2. TP53 in particular is anti-correlated with most other events.

Although cMHN and oMHN both fit the data similarly well, they offer drastically different causal interpretations (Fig. 4): cMHN suggests that APC, KRAS and TP53 strongly antagonize each other as well as other events. oMHN instead proposes that APC, KRAS and especially TP53 lead to observation. This explains away many of their suppressive interactions and even uncovers a synergy between APC und TP53.

**Fig. 4.**
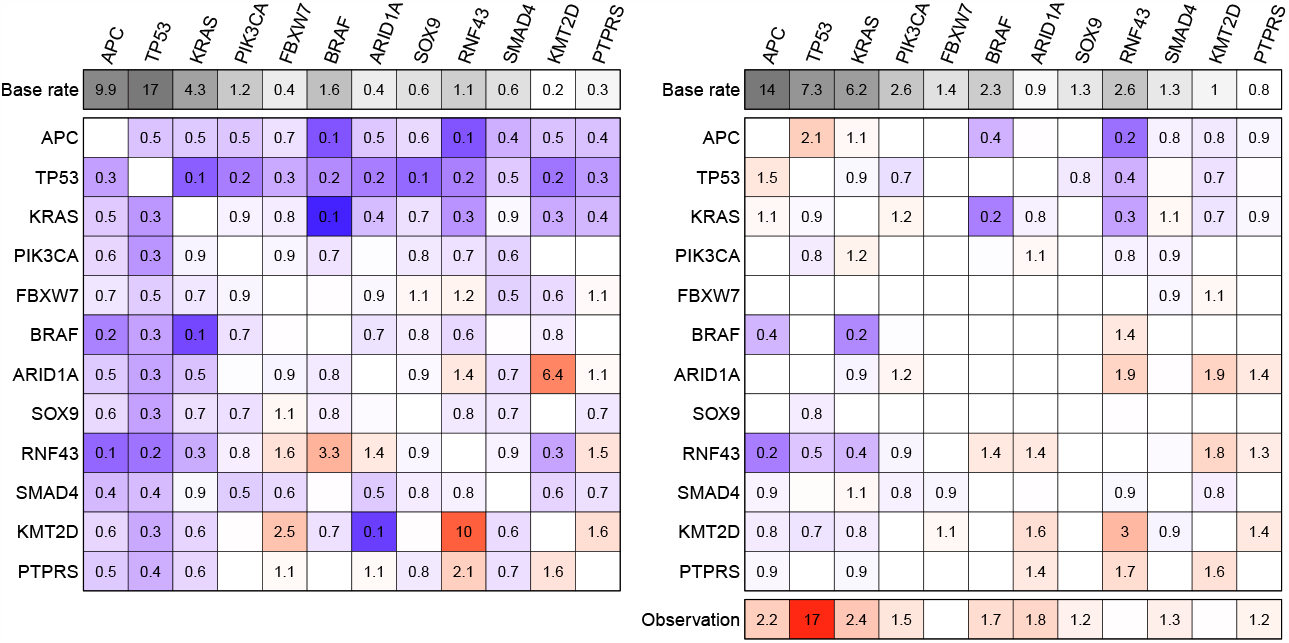
Heatmap visualization of the cMHN (left) and the oMHN (right) for the COAD dataset. In the main heatmap bodies, each cell shows the multiplicative effect *Θ*_*ij*_ from the column event *j* on the row event *i*. Promoting effects > 1 are coded in red, suppressive effects *<* 1 in blue and neutral effects = 1 are blank. The additional top row indicates base rates *Θ*_*ii*_ and the bottom row indicates effects Ω_*j*_ from each column event *j* on the observation event. Values are rounded to the first decimal.

The different causal models also imply different chronological orders of events. For a given tumor genotype, we consider the probability of every possible chronological order according to each model, see Table 1. The most probable orders for common genotypes in the dataset are shown in Fig. 5. Contrary to cMHN, the oMHN suggests that APC tends to occur early in the progression and that TP53 tends to occur late, despite its prevalence. This is because TP53 triggers and therefore immediately precedes the observation.

**Table 1.** All possible chronological orders to reach the genotype that contains exactly the events APC, KRAS, TP53 and was observed. The probabilities of these orders according to cMHN and oMHN are computed as in Supplementary S3 and rounded to the 3rd decimal.

| Chronological order | cMHN | oMHN |
| --- | --- | --- |
| APC → KRAS → TP53 | 0.068 | 0.312 |
| KRAS → APC → TP53 | 0.064 | 0.262 |
| APC → TP53 → KRAS | 0.198 | 0.149 |
| KRAS → TP53 → APC | 0.075 | 0.113 |
| TP53 → APC → KRAS | 0.383 | 0.101 |
| TP53 → KRAS → APC | 0.212 | 0.064 |

**Fig. 5.**
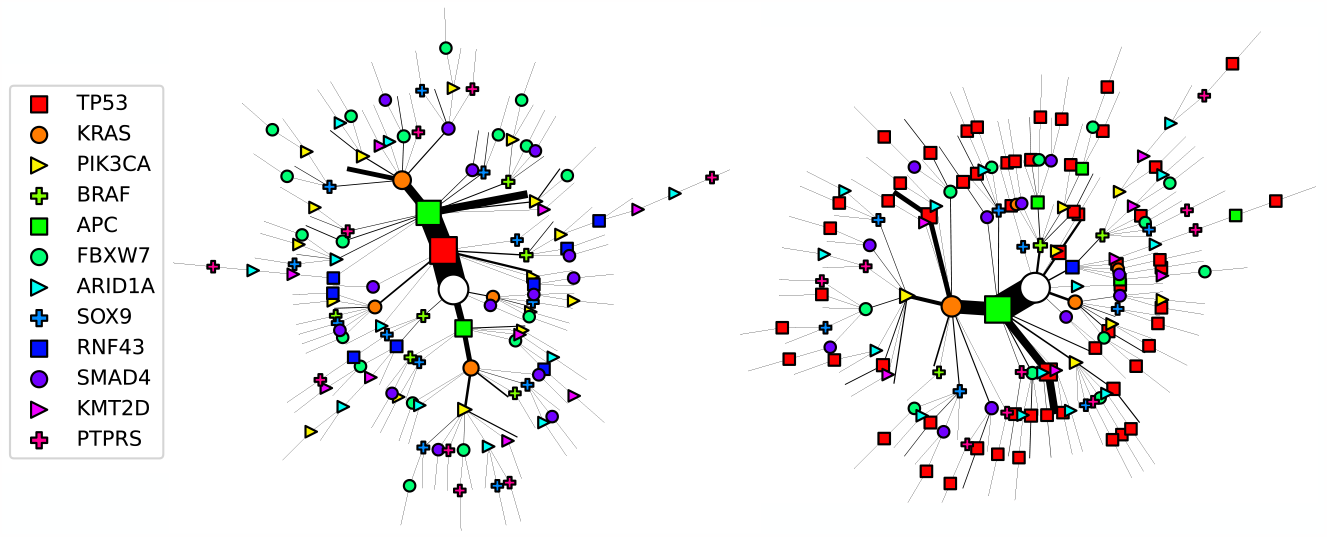
Most probable chronological order of events for the COAD dataset according to the cMHN (left) and oMHN (right). Each path from the root of the tree (white circle) to a leaf represents the progression of a tumor in the dataset. The symbols along the path indicate events whose most probable chronological order was computed from the trained models. To avoid clutter, the observation event at every leaf is implied without drawing a symbol and only tumors whose state is shared by at least 3 patients are drawn. The size of the edges and symbols along a path scale in the total number of patients with that tumor state.

Unlike for cMHN, the interpretations drawn from oMHN are in line with common conceptions about COAD genetic progression. It has been repeatedly suggested that APC inactivation is a gatekeeper which starts the transformation in healthy tissues and is a prerequisite for subsequent alterations, like TP53, which then elicit aggressive growth and invasion [10,53,55]. The synergy between APC and TP53 is further supported by a systematic study of conditional selection effects in cancer genomes [29] which found that TP53 mutations are under particularly strong positive selection in APC-mutated colorectal cancers, and vice versa. However, oMHN also suggests that TP53 alterations are able to generate clinically conspicuous tumors on their own.

Despite these differences, some interactions remain consistent between cMHN and oMHN. Most notably, both models suggest a “double antagonism” between the KRAS-APC and BRAF-RNF43 pairs: each event in one pair suppresses both members of the other pair. In fact, the two event pairs likely produce similar consequences through alternative means: both event pairs deregulate the RAS and Wnt pathways. These are synergistic milestones in COAD progression [30,33]. Specifically, KRAS and BRAF mutations are alternative ways of RAS pathway deregulation [13,44] and APC and RNF43 are alternative ways of Wnt signalling deregulation [21,23]. Additionally, the synergy within the pairs as well as the antagonism between them are clearly reflected in the conditional selection analysis of [29]. Both points, functional similarity and conditional selection effects, support genuine antagonism between these pairs.

Interestingly, the BRAF-RNF43 pair is associated with a distinct mode of COAD progression, the Serrated Neoplasia Pathway. These cancers develop from serrated sessile lesions, with different histopathological and prognostic properties [34]. Unlike in conventional COADs, APC mutations are rare here while BRAF mutations are thought to be initial [8,6]. Experimental evidence suggests that specifically MLH1-deficient, microsatellite-instable serrated COADs rely on BRAF and RNF43 mutations in their progression [7,54].

Taken together, these findings suggest that there are two prototypical ways of genetic progression in COAD. On the one hand, any combination of the synergistic triplet APC-KRAS-TP53 can be sufficient to elicit observation, although APC tends to be the initiating factor and TP53 the observation driver. On the other hand, crucial pathway deregulation can also be achieved by alternatives like BRAF and RNF43. In these cases, there is no main observation driver and typically more alterations are accumulated before observation.

### 3.2 Lung Adenocarcinoma

In the models on lung adenocarcinoma (LUAD), Fig. 6, we also observed a shift from widespread suppressive interactions in cMHN to obervation rate increases in oMHN, most notably for EGFR mutations. EGFR mutations appear mutually exclusive with many other events in the input data. cMHN models this with widespread suppressive interactions while oMHN explains these away by an observation rate increase. For EGFR and TP53, oMHN even suggests synergy instead of the antagonism proposed by cMHN.

**Fig. 6.**
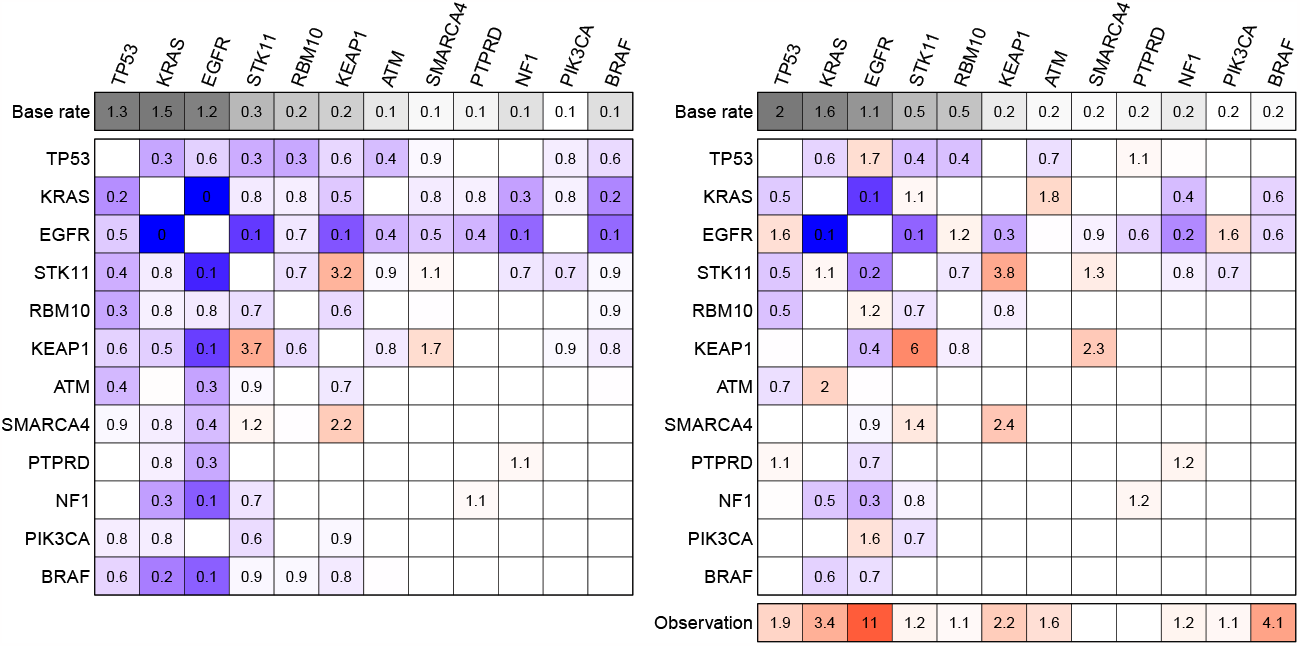
Heatmap visualization of the cMHN (left) and the oMHN (right) for the LUAD dataset. In the main heatmap bodies, each cell shows the multiplicative effect *Θ*_*ij*_ from the column event *j* on the row event *i*. Promoting effects > 1 are coded in red, suppressive effects *<* 1 in blue and neutral effects = 1 are blank. The additional top row indicates base rates *Θ*_*ii*_ and the bottom row indicates effects Ω_*j*_ from each column event *j* on the observation event. Values are rounded to the first decimal.

The chronological orders differ between cMHN and oMHN mainly for EGFR and KRAS. These events tend to occur later according to oMHN because they trigger observation, see Fig. 7. According to oMHN, EGFR has a strong effect on observation on its own. Conversely, KRAS-positive LUADs elicit observation in a more concerted manner supported by e.g., ATM, STK11 and KEAP1. Moreover, there are also interactions that remain consistent between cMHN and oMHN, most prominently the suppressive relationship between EGFR and KRAS. In fact, experimental demonstration of synthetic lethality [52] and conditional selection analysis [29] both support a genuine antagonism.

**Fig. 7.**
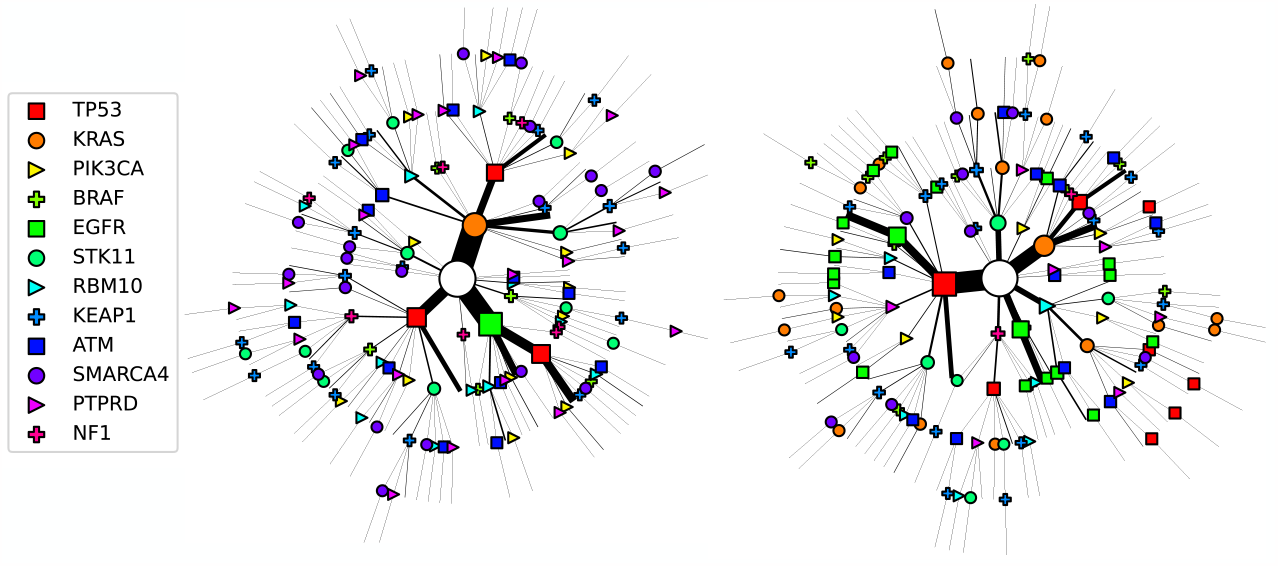
Most probable chronological orders of events for the LUAD dataset according to the cMHN (left) and oMHN (right). Each path from the root of the tree (white circle) to a leaf represents the progression of a tumor in the dataset. The symbols along the path indicate events whose most probable chronological order was computed from the trained models. To avoid clutter, the observation event at every leaf is implied without drawing a symbol and only tumors whose state is shared by at least 3 patients are drawn. The size of the edges and symbols along a path scale in the total number of patients with that tumor state.

## 4 Discussion

Large cancer genomics datasets offer a valuable opportunity for modeling cancer progression, but many of them are observational, drawn from routine clinical practice rather than controlled trials [51]. This makes them prone to pervasive biases [25], such as the notorious confounder bias, which is due to unaccounted effects from latent variables, and the collider bias, which is due to unaccounted effects on a conditioned outcome. In this paper, we have resolved an important instance of the collider bias by learning which genetic events cause the clinical observation of a tumor. This is an important biological insight on its own, since the observation of a tumor is often tied to its size and aggressiveness. In addition, and perhaps more consequentially, learning their effects explains away spurious interactions between many other events and uncovers interactions that were previously hidden.

While resolving the collider bias is a crucial step towards more reliable cancer progression models, other sources of confounding may still remain. Future work should combine our approach with the modeling of latent variables such as environmental factors, mutational processes, or the tumor’s cell type of origin. Further non-identifiability could be mitigated by exploiting known time intervals between consecutive observations [48], such as biopsies of primary tumors and metastases. Surprisingly, [22] have shown that the identifiability of classical MHNs can also be improved simply by including more events, which becomes possible through efficient learning algorithms [32,19,22]. Moreover, our approach could also be applied to models of subclonal tumor compositions [36].

More realistic models will ultimately allow us to not only understand and predict the course of cancer progression, but also to steer it towards favorable outcomes through treatment.

## 5. Acknowledgements

This work was supported by the Swiss National Science Foundation grant 179518, the Swiss Cancer League grant KFS-2977-08-2012 and the German Research Foundation grants TRR-305 and GR-3179/6-1.

## Footnotes

4 This construction is only needed for learning the model. In order to extrapolate the progression of a tumor into the future beyond its observation, one would “unfreeze” the process again by setting the outgoing effects of the observation to 1. Ideally one would include effects from the treatment of the patient instead.

5 In the independence model, events occur independently of each other with rates equal to their odds in the dataset.

