## Supplement for "Overcoming Observation Bias for Cancer Progression Modeling"

[1] Department of Biosystems Science and Engineering, ETH Zürich, Basel, Switzerland
{Rudolf.Schill,Niko.Beerenwinkel}@bsse.ethz.ch
[2] Department of Statistical Bioinformatics, University of Regensburg, Germany
[3] Institute for Geometry and Applied Mathematics, RWTH Aachen, Germany

### S1 Data acquisition and preprocessing

We obtained the GENIE release 13.1-public dataset through synapse.org [6]. We selected entries from MSKCC's MSK-IMPACT project [3] and further filtered for primary tumor samples with COAD or LUAD Oncotree codes, respectively. If multiple primary tumor samples were available per patient, we selected only the sample with the lowest age at sequencing. This yielded 2269 samples for COAD and 3662 samples for LUAD, respectively. We additionally annotated mutation data with OncoKB [2]. To restrict the data to likely functional variants, we carried out the following filtering scheme:

1. If OncoKB annotation for oncogenicity was present, accept variants that are annotated "Oncogenic", "Likely Oncogenic" or "Resistance" and omit variants that are annotated "Neutral" or "Likely Neutral". If the annotation is not present, proceed with 2.
2. If either PolyPhen-2 [1] or SIFT [5] annotations were present, accept variants that are annotated "Probably damaging" (PolyPhen-2) and/or "deleterious" (SIFT). Accept variants that are annotated "Possibly damaging" (PolyPhen-2) and "deleterious (low conf)"/no annotation (SIFT). Accept variants that have no PolyPhen-2 annotation but are annotated "deleterious (low conf)" (SIFT). Omit all other variants that have either annotation. If no annotation is present, proceed with 3.
3. Variant Class annotations were present for all variants. Accept variants that are annotated "Frame shift Ins/Del", "Nonsense mutation", "Nonstop mutation", "Splice site" and "Translation start site". Omit variants that are annotated "In-frame Ins/Del", "Misssense mutation", "3'/5'-Flank", "3'/5'-UTR", "Intron", "Splice region", "Silent".

Furthermore, we restricted our analysis to the 341 genes that are assayed by MSK-IMPACT since its earliest version [3]. Within genes, we made no distinction between types of functional variants. We restricted our analysis to the 12 most commonly affected genes per cancer type.

### S2 Event load enrichment testing

To ascertain the enrichment of certain events in samples with few or many active events, we compared the observed distribution with randomized distributions via Z-scores (see Fig. S1). Specifically, to randomize a given dataset, we assigned each sample a random count of active events ("event load", drawn from the original distribution) and as many random active events (drawn from the original distribution). To calculate Z-scores, we randomized datasets 10000 times for COAD and LUAD each.

| | APC | TP53 | KRAS | PIK3CA | FBXW7 | BRAF | ARID1A | SOX9 | RNF43 | SMAD4 | KMT2D | PTPRS |
|---|---|---|---|---|---|---|---|---|---|---|---|---|
| L1 | 0.4 | 9.9 | −0.2 | −3.9 | −3.1 | −1.6 | −2.9 | −2.2 | −2.2 | −1.7 | −2.4 | −1.7 |
| L2 | 10.2 | 12.1 | −2.7 | −7.0 | −6.4 | −1.5 | −6.1 | −4.9 | −5.0 | −4.0 | −5.9 | −6.0 |
| L3 | 3.7 | 3.5 | 5.8 | −0.0 | −3.7 | −1.9 | −5.0 | −3.2 | −4.7 | 0.0 | −6.2 | −4.4 |
| L4 | −1.9 | −4.4 | 1.5 | 3.4 | 2.7 | 1.0 | 1.2 | 2.6 | 0.2 | 2.5 | −0.2 | −0.8 |
| L5 | −4.2 | −5.7 | −1.3 | 2.7 | 4.3 | 0.4 | 4.6 | 3.0 | 3.6 | 1.7 | 4.9 | 5.3 |
| L6 | −4.4 | −5.5 | −3.4 | 1.6 | 3.1 | 2.4 | 5.0 | 3.4 | 6.8 | 0.8 | 7.4 | 6.5 |
| L7 | −3.9 | −3.2 | −3.1 | 0.5 | 3.3 | 2.2 | 5.8 | 1.7 | 5.0 | −0.7 | 6.6 | 4.5 |
| L8 | −1.4 | −1.6 | −0.4 | 0.7 | 2.8 | −0.9 | 2.4 | 1.0 | −0.3 | −0.7 | 3.2 | 2.7 |
| L9 | −0.4 | −0.4 | −0.8 | 0.3 | 0.7 | −0.5 | 0.9 | 0.9 | 0.9 | −0.5 | 1.0 | 1.1 |
| L10 | −0.7 | −0.7 | −0.7 | 0.3 | 0.9 | 1.0 | 0.1 | 0.1 | 0.2 | 1.1 | 1.2 | 1.4 |

| | TP53 | KRAS | EGFR | STK11 | RBM10 | KEAP1 | ATM | SMARCA4 | PTPRD | NF1 | PIK3CA | BRAF |
|---|---|---|---|---|---|---|---|---|---|---|---|---|
| L1 | −1.5 | 9.5 | 13.4 | −6.6 | −5.1 | −7.5 | −4.6 | −5.4 | −4.7 | −3.0 | −4.7 | 2.4 |
| L2 | 7.1 | −2.2 | 5.4 | −3.6 | −0.0 | −4.4 | −1.7 | −4.6 | −4.5 | −1.2 | −0.4 | −2.0 |
| L3 | −3.0 | −1.4 | −8.7 | 5.0 | 2.6 | 5.6 | 3.9 | 3.1 | 3.4 | 0.4 | 3.2 | −0.0 |
| L4 | −4.0 | −2.3 | −7.7 | 5.6 | 1.1 | 7.1 | 1.7 | 5.9 | 6.5 | 3.2 | 0.5 | 1.0 |
| L5 | −2.4 | −1.8 | −3.4 | 0.8 | 1.3 | 1.6 | 1.7 | 6.6 | 4.1 | 3.4 | 2.0 | −0.5 |
| L6 | −1.0 | −1.2 | −1.6 | 1.3 | 0.6 | 2.0 | −0.0 | 3.6 | 0.9 | 0.9 | 0.1 | 0.2 |

**Fig. S1.** Event load enrichment test results for COAD (left) and LUAD (right). Rows correspond to event loads (e.g., L2: samples with exactly two active events) and columns correspond to events. Each cell shows the corresponding Z-score.

### S3 Computing Order Probabilities

To compute the probability of an order of events according to oMHN, we use a formula inspired by [4, eq.s 4 and 6]:

Let $\sigma = (\sigma_1, \dots, \sigma_k)$ be a tuple of $k$ events. For $i \in \{1, \dots, k\}$ let $\sigma_{[:i]} \coloneqq \{\sigma_1, \dots, \sigma_i\}$. Then

$$\begin{aligned}
\mathbf{P}(\text{Observation of } \sigma) &= \left( \prod_{i=1}^{k} \mathbf{P}\Big(\text{first } i \text{ events are } \sigma_{[:i]} \mid \text{first } i-1 \text{ events are } \sigma_{[:i-1]}\Big) \right) \\
&\quad \cdot \mathbf{P}\Big(\text{Observation of } \sigma \mid \text{first } k \text{ events are } \sigma_{[:k]}\Big) \\
&= \left( \prod_{i=1}^{k} \frac{Q_{\sigma_{[:i-1]}, \sigma_{[:i]}}}{\prod_{j \in \sigma_{[:i-1]}} \Omega_j + \sum_{j \notin \sigma_{[:i-1]}} Q_{\sigma_{[:i-1]}, \sigma_{[:i-1]+j}}} \right) \frac{\prod_{j \in \sigma_{[:k]}} \Omega_j}{\prod_{j \in \sigma_{[:k]}} \Omega_j + \sum_{j \notin \sigma_{[:k]}} Q_{\sigma_{[:k]}, \sigma_{[:k]+j}}} \\
&= \left( \prod_{i=1}^{k} \frac{Q_{\sigma_{[:i-1]}, \sigma_{[:i]}}}{\prod_{j \in \sigma_{[:i-1]}} \Omega_j - Q_{\sigma_{[:i-1]}, \sigma_{[:i-1]}}} \right) \frac{\prod_{j \in \sigma_{[:k]}} \Omega_j}{\prod_{j \in \sigma_{[:k]}} \Omega_j - Q_{\sigma_{[:k]}, \sigma_{[:k]}}} \\
&= \left( \prod_{i=1}^{k} \frac{\hat{Q}_{\sigma_{[:i-1]}, \sigma_{[:i]}}}{1 - \hat{Q}_{\sigma_{[:i-1]}, \sigma_{[:i-1]}}} \right) \frac{1}{1 - \hat{Q}_{\sigma_{[:k]}, \sigma_{[:k]}}},
\end{aligned}$$

where $\hat{Q}$ is the matrix from eq. (15)

$$\hat{Q} = \sum_{i=1}^{n} \bigotimes_{j=1}^{i-1} \begin{pmatrix} 1 & 0 \\ 0 & \Theta_{ij}/\Omega_j \end{pmatrix} \otimes \begin{pmatrix} -\Theta_{ii} & 0 \\ \Theta_{ii} & 0 \end{pmatrix} \otimes \bigotimes_{j=i+1}^{n} \begin{pmatrix} 1 & 0 \\ 0 & \Theta_{ij}/\Omega_j \end{pmatrix} .$$
